# Epigenetic control of myogenic identity of human muscle stem cells in Duchenne Muscular Dystrophy

**DOI:** 10.1101/2023.04.26.538414

**Authors:** Jimmy Massenet, Michèle Weiss-Gayet, Hina Bandukwala, Mélanie Magnan, Arnaud Hubas, Patrick Nusbaum, Isabelle Desguerre, Cyril Gitiaux, F Jeffrey Dilworth, Bénédicte Chazaud

## Abstract

In Duchenne Muscular Dystrophy (DMD), the absence of the subsarcolemmal dystrophin protein leads to repeated myofiber damages inducing cycles of muscle regeneration that is driven by muscle stem cells (MuSCs). With time, MuSC regenerative capacities are overwhelmed, leading to fibrosis and muscle atrophy. Whether MuSCs from DMD muscle have intrinsic defects that limit regenerative potential or are disrupted by their degenerative/regenerative environment is unclear. We investigated cell behavior and gene expression in human using MuSCs derived from DMD or healthy muscles. We found that proliferation, differentiation and fusion were not altered in DMD-MuSCs, but with time, they lost their myogenic identity twice as fast as healthy MuSCs. The rapid drift towards a fibroblast-like cell identity was observed at the clonal level, and resulted from the altered expression of epigenetic enzymes required to maintain the myogenic cell fate. Indeed, the re-expression of *CBX3*, *SMC3*, *H2AFV* and *H3F3B* prevented the MuSC identity drift. Amongst the epigenetic changes, a closing of chromatin at the gene encoding the transcription factor *MEF2B* caused a down-regulation of its expression and a loss of the myogenic fate. Thus, MEF2B is a key mediator of the myogenic identity in human MuSCs, that is altered in DMD pathology.

## Introduction

After an injury, adult skeletal muscle regenerates thanks to muscle stem cells (MuSCs), or satellite cells, that operate adult myogenesis to ensure the formation of new myofibers while repopulating the pool of stem cells for further needs (Wosczyna and Rando 2018). To execute myogenesis, MuSCs transit through sequential states including activation (exit from quiescence), proliferation (expansion), exit from the cell cycle and commitment into terminal myogenic differentiation, and eventually fusion into multinucleated myotubes and myofibers (Wosczyna and Rando 2018). The characteristics of each cell fate are determined through specific epigenetic mechanisms that determine the subset of genes that are expressed, through the control of the accessibility of the transcriptional machinery to specific loci. The chromatin state allows the expression of the ad hoc genes in a spacio-temporal manner. Changes in chromatin organization is required for mediating decisions of cell fate and differentiation (Massenet et al. 2021). In response to environmental cues, chromatin organization is altered and modifies the accessibility of the transcription machinery to the gene sequences, which is modulated by several levels of regulation, at the DNA and histone levels. Huge efforts have been made to decipher the chromatin organization and the key epigenetic regulators that control adult myogenesis (Aziz et al. 2010; Asp et al. 2011; Segalés et al. 2015). Investigations were mainly done in the context of regeneration of healthy muscle.

Epigenetic marks and chromatin dynamics are reversible and change according to the modifications of the environment of the cells. Such environmental changes are particularly important during muscular diseases, and particularly during degenerative myopathies, where attempts of muscle regeneration occur in an environment encompassing tissue damage, inflammation and fibrosis. However, chromatin dynamic response to environmental changes and how it impacts of muscle regeneration has been poorly investigated in the context of muscular diseases. Duchenne Muscular dystrophy (DMD) is caused by mutations in the dystrophin gene (Hoffman et al. 1987; Dalkilic and Kunkel 2003; Mendell and Lloyd-Puryear 2013). The absence of dystrophin causes sarcolemma fragility and costamere disorganization, leading to myofiber damage (Petrof et al. 1993; Williams and Bloch 1999). Patient muscles present asynchronous cycles of damage and regeneration and are characterized by a progressive loss of muscle tissue associated with chronic inflammation and fibrosis. The defect in muscle repair that is finally observed in DMD patients has been attributed to both MuSC cell-autonomous and non-autonomous mechanisms. Most of the investigations were done in the mdx mouse model of DMD, which poorly recapitulates the clinical features observed in DMD patient’s muscle (Partridge 2013; Bareja et al. 2014). Studies reported intrinsic alterations of MuSC differentiation and self-renewal capacities (Yablonka-Reuveni and Anderson 2006; Dumont et al. 2015a) while others reported that the MuSC environment directly impacts on MuSC function (Boldrin et al. 2015). Finally, lineage tracing experiments reported that in the mdx muscle, a portion (7-20%) of MuSCs acquires a fibroblastic phenotype, suggesting strong alteration of chromatin organization in such converted cells in response to environmental cues (Brack et al. 2007; Biressi et al. 2014; Pessina et al. 2015).

Given the difficulties to unravel the mechanisms of failure of muscle repair in DMD in the mdx model, using human DMD MuSCs is an attractive alternative. Pioneer investigations using cells isolated from human muscle biopsies led to contradictory results about the myogenic potential of cells issued from DMD muscle as compared with cells isolated from healthy muscle (Franklin et al. 1981; Ionasescu and Ionasescu 1982; Blau et al. 1983a; Blau et al. 1983b; Jasmin et al. 1984; Iannaccone et al. 1987). Discrepant results are likely due to the lack of efficient isolation procedure of human MuSCs at that time, leading to the analysis of mixed cultures containing various cell types. During the early 2000’s, isolation techniques were developed, using FACs or magnetic cell sorting, allowing the obtention of highly purified (more than 95-98%) MuSCs. Those techniques were based on the expression of CD56 (or Neural Cell Adhesion Molecule, NCAM) by MuSCs, expression which was established as a reliable marker of myogenicity of human MuSCs (Illa et al. 1992; Stewart et al. 2003; Mackey et al. 2009; Agley et al. 2013; Xu et al. 2015; Uezumi et al. 2016).

In the present study, we examined the myogenic potential and the myogenic identity of human MuSCs isolated from DMD muscle as compared with healthy MuSCs. Being isolated at the time of diagnosis, DMD-MuSCs have been supposedly living over time with the constant presence of stressors around, that may impact they chromatin organization. We functionally investigated some features of chromatin dynamics in DMD-MuSCs, that may explain the rapid identity drift observed in these cells and we identified epigenetic regulators involved in the maintenance of the myogenic identity of human MuSCs.

## Results

Human Muscle stem cells (MuSCs) were obtained from the hospital cell bank as more than 98% of CD56^pos^ cells (cells were previously expanded and sorted based on their CD56 expression [Fig. S1A,B]). CD56 was established as a reliable marker of myogenicity of human MuSCs (Illa et al. 1992; Stewart et al. 2003; Mackey et al. 2009; Agley et al. 2013; Xu et al. 2015; Uezumi et al. 2016). Indeed, 100% of CD56^pos^ cells expressed the transcription factor Pax7 (Fig. S1C).

### CD56^pos^ DMD-MuSCs exhibit normal myogenic properties

CD56^pos^ MuSCs derived from DMD (DMD-MuSCs) and healthy control (HC-MuSCs) muscles were cultured to evaluate their capacity to perform *in vitro* myogenesis. Proliferation of CD56^pos^ cells cultured in growth medium, assessed by EdU incorporation, was similar in HC- and DMD-MuSCs (Fig. 1A). Commitment into terminal myogenic differentiation of CD56^pos^ cells, assessed by their expression of myogenin when cultured in differentiation medium, was also not different in HC- and DMD-MuSCs (Fig. 1B). Finally, a fusion assay of differentiated cells (myocytes) showed similar capacity of HC- and DMD-MuSCs to form myotubes (Fig. 1C). These data show that the myogenic capacities were not altered in CD56^pos^ DMD-MuSCs.

**Fig. 1.**
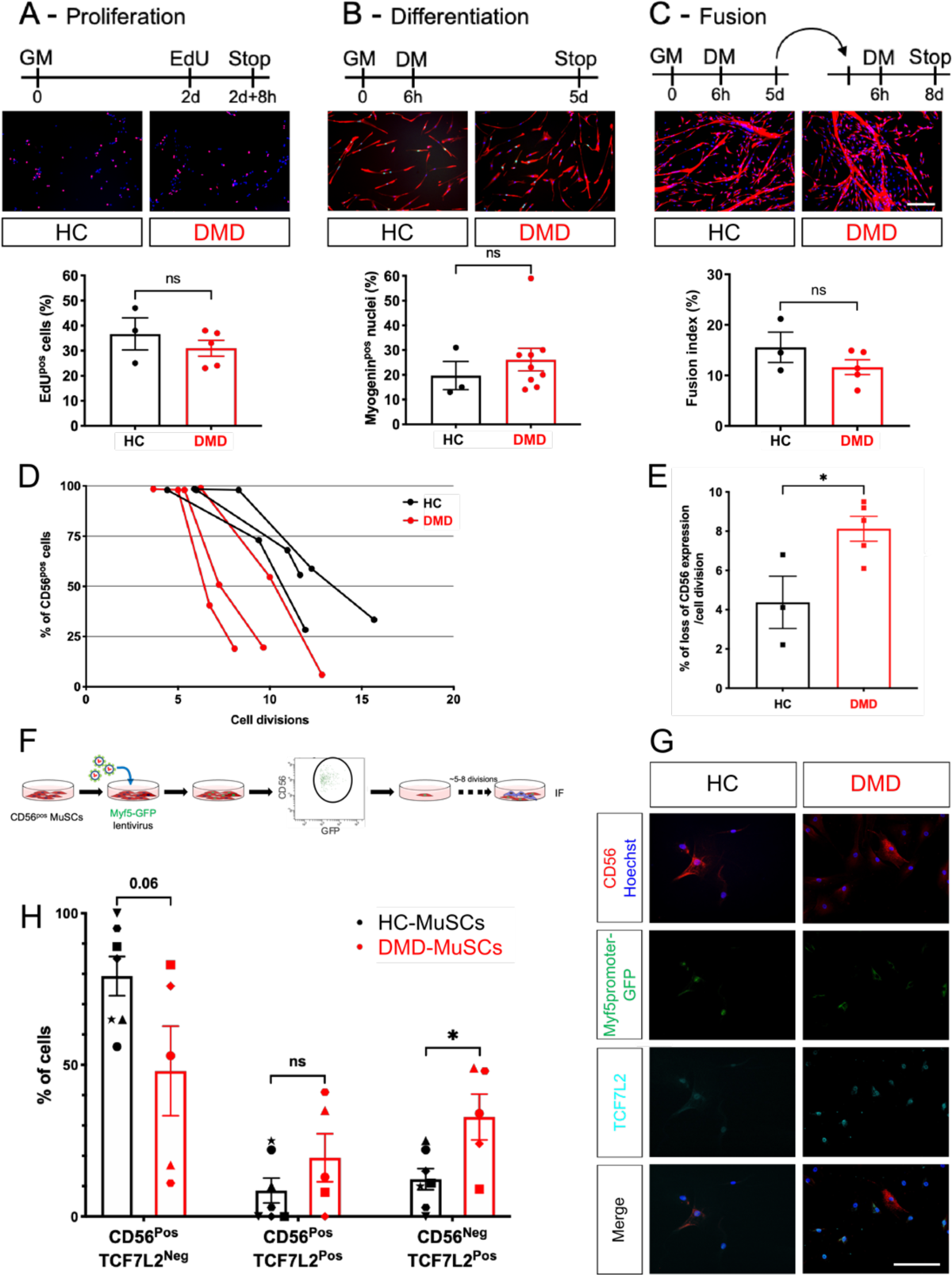
*In vitro* behavior of DMD-MuSCs. **(A-C)** CD56^pos^ MuSCs isolated from healthy control (HC) and Duchenne (DMD) muscles were analyzed for their capacity to implement *in vitro* myogenesis. **(A)** Proliferation was assessed in growth medium (GM) as the number of EdU^pos^ cells (red). **(B)** Differentiation was quantified after 5 days in differentiation medium (DM) as the number of myogenin^pos^ cells (green) among desmin^pos^ cells (red). **(C)** Fusion index was quantified in differentiated cells grown at high density, as the number of nuclei in desmin expressing myotubes (red) related to the total number of nuclei. Hoechst labels nuclei (blue). Bar = 100 μm. **(D)** Expression of CD56 was evaluated by flow cytometry during the culture of initially pure CD56^pos^ HC- and DMD-MuSCs in growth medium. **(E)** Calculation of the loss of CD56 per cell division from (D). **(F)** Experimental procedure of the clonal culture of Myf5-transduced CD56^pos^ MuSCs. **(G)** Immunostaining of clones for CD56 (red), GFP(Myf5) (green) and TCF7L2 (cyan). Hoechst labels nuclei (blue). Bar = 50 μm. **(H)** Quantification of cells according to the immunostaining shown in (G). Each shape symbol represents one clone. Results are means ± SEM of 3 to 9 samples in A-E and of 6 clones issued from 3 HC donors and of 5 clones issued from 3 DMD donors in H. ns: non significant, *p<0.05 using unpaired (A-E) or paired (H) t-test.

### DMD-MuSCs lose their myogenic identity twice as fast as HC-MuSCs

A progressive loss of CD56 expression has been described during the culture of human CD56^pos^ MuSCs issued from healthy muscle, preventing their use after 4-5 passages (Stewart et al. 2003; Agley et al. 2013; Alsharidah et al. 2013; Kim et al. 2021). Starting from 100% CD56^pos^ HC- and DMD-MuSC populations, we found that the number of CD56^pos^ cells decreased with time in culture, as expected, however this decrease occurred earlier and faster in DMD-MuSCs (Fig. 1D). The rate of CD56^pos^ cells lost per cell division was calculated over a period of 10 population doublings and was found to be twice as high in DMD-MuSCs *vs.* HC-MuSCs (8.1±0.6% *vs*. 4.4±1.3% per cell division) (Fig. 1E). The loss of CD56^pos^ cells could be due to either apoptosis of CD56^pos^ cells, overtake of CD56^neg^ cells in the culture (although being not more than 2% of the cells in the starting cultures), or cellular conversion of CD56^pos^ into CD56^neg^ cells. To investigate apoptosis, CD56^pos^ MuSCs were maintained in growth medium until about half of the cells have lost CD56 expression, then cells were sorted as CD56^pos^ and CD56^neg^ cell populations (Fig. 2A) and were analyzed. TUNEL assay indicated that both CD56^pos^ and CD56^neg^ cells from HC and DMD donors showed very low rates of apoptosis (from 1.7 to 3.3% of the cultured cells) (Fig. S2A), ruling out apoptosis as a mechanism for CD56 loss in MuSC population. We then used cells from adult healthy samples to compare the growth rate and CD56 expression of 100% CD56^pos^ cells, 100% CD56^neg^ cells and a mixed culture of 50:50 CD56^pos^:CD56^neg^ cells. Both growth rate and loss of CD56 expression did not differ between the 3 conditions (Fig. S2B,C), indicating that the presence of CD56^neg^ cells in the culture did not impact on CD56^pos^ cell behavior, and ruling out a faster expansion of CD56^neg^ cells to explain the CD56 loss in MuSC cultures.

**Fig. 2.**
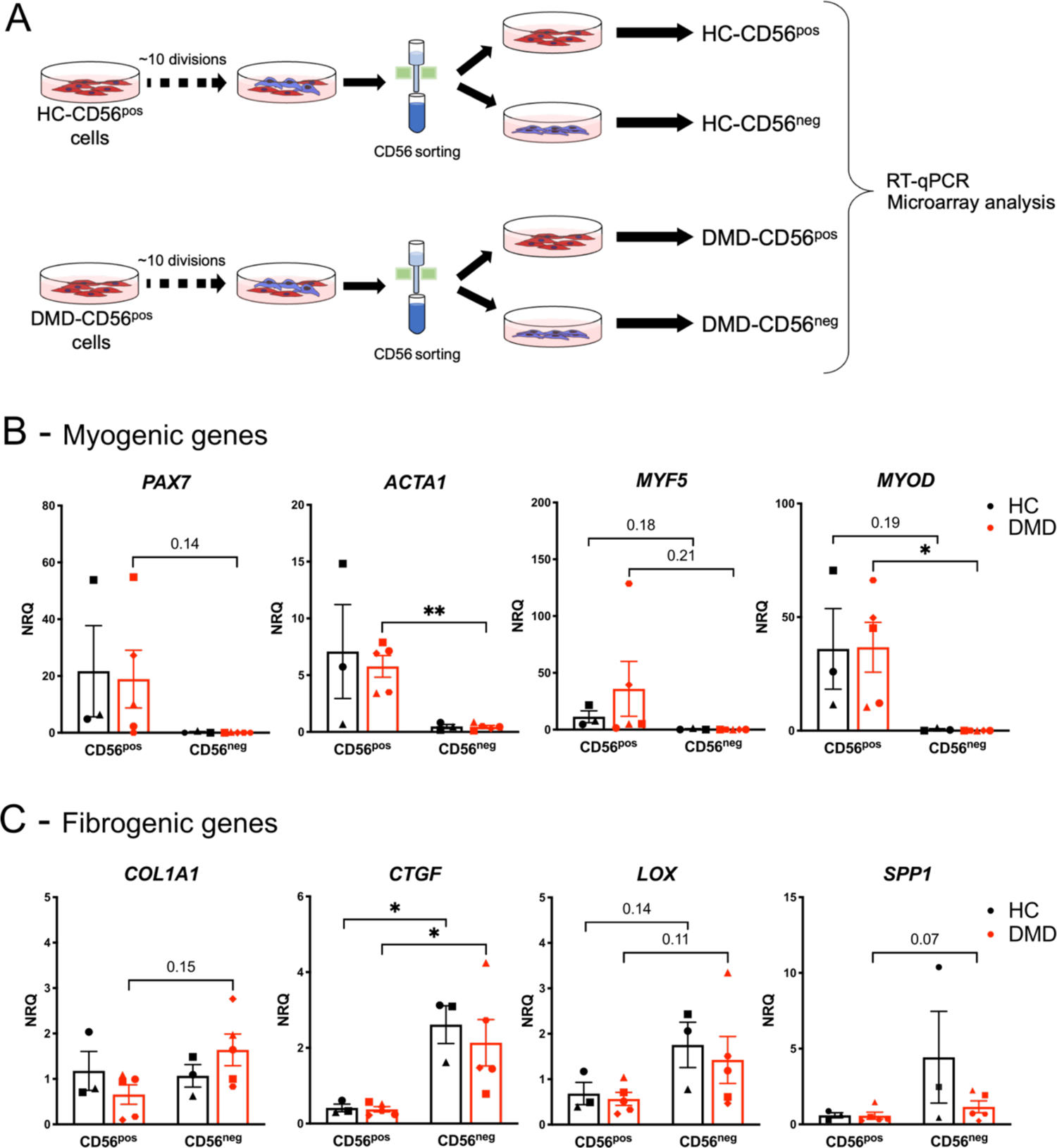
Gene expression in CD56^pos^ and CD56^neg^ cells. **(A)** Experimental procedure for CD56^pos^ and CD56^neg^ cell purification originating from pure healthy control (HC)- and Duchenne (DMD) CD56^pos^ population. Cells were cultured in growth medium. **(B-C)** Normalized Relative Quantity (NRQ) expression by CD56^pos^ and CD56^neg^ cells from HC- and DMD-samples evaluated by RT-qPCR of **(B)** the myogenic related genes *PAX7, ACTA1, MYF5* and *MYOD* and **(C)** the fibrogenic related genes *COL1A1*, *CTGF*, *LOX* and *SPP1*. Results are means ± SEM of 3 to 6 samples. Each shape symbol represents cells issued from one initial culture. *p<0.05, **p<0.001 using paired t-test.

To futher investigate the conversion of CD56^pos^ cells into CD56^neg^ cells, we performed clonal cell cultures. CD56^pos^ cells from HC and DMD donors were transduced with lentiviruses encoding GFP under the control of the *Myf5* promoter region. Double CD56^pos^/GFP^pos^ cells were single cell sorted and seeded in 96-well plates (Fig. 1F). Examination of each well confirmed the presence of only a single CD56^pos^/GFP^pos^ cell (Fig. S1D). Clones were grown for 4 to 6 weeks in proliferating medium, then immunostained for CD56 and TCF7L2, a transcription factor expressed by human muscle fibroblasts and fibroadipogenic precursors (Mackey et al. 2017; Farup et al. 2021) (Fig. 1G). Clones derived from DMD-MuSCs gave rise to 40% less CD56^pos^/TCF7L2^neg^ cells than those derived from HC-MuSCs (48 *vs.* 79%) and they produced 2.6-fold more CD56^neg^/TCF7L2^pos^ cells (33 *vs.* 13%) (Fig. 1H). A small proportion of double positive cells was counted in both HC and DMD cultures, possibly representing an intermediate status (Fig. 1H). These results show at the cellular level the conversion of CD56^pos^ MuSCs into CD56^neg^/TCF7L2^pos^ cells, which was higher in DMD than in HC cultures.

To analyze the nature of the cells at the transcriptomic level, CD56^pos^ HC- and DMD-MuSCs were cultured until about 50% of the cells have lost CD56 expression, and cells were further sorted to obtained CD56^pos^ and CD56^neg^ populations issued from the same initial MuSCs (hereafter referred as HC-CD56^pos^, HC-CD56^neg^, DMD-CD56^pos^ and DMD-CD56^neg^) (Fig. 2A). RT-qPCR experiments showed a dramatic decrease of the expression of the muscle-specific genes *PAX7, MYOD, MYF5* and *ACTA1* genes in both HC- and DMD-CD56^neg^ cells, confirming the loss of myogenicity of these cells (Fig. 2B). Inversely, the expression of genes associated with fibrogenesis, *COL1A1*, *CTGF*, *LOX* and *SPP1*, was increased in CD56^neg^ cells (Fig. 2C). No difference was observed between HC- and DMD-derived cells. Altogether these results show that MuSCs from DMD muscle lose their myogenicity to acquire fibrogenic-like features faster than cells issued from normal muscle.

### Reduced expression of epigenetic regulatory factors accompanies the loss of myogenicity in HC-CD56^neg^ cells and characterizes DMD-CD56^pos^ cells

We performed transcriptomic analysis on the four populations (HC-CD56^pos^, HC-CD56^neg^, DMD-CD56^pos^ and DMD-CD56^neg^ as described in Fig. 2A) using several comparisons. Gene ontology analysis of differentially expressed genes (Table S1) between CD56^neg^ and CD56^pos^ cells in both HC and DMD samples showed a common downregulation of genes involved in muscle function and an overexpression of genes involved in extracellular matrix (ECM) (Fig. S3, “common in HC and DMD”), in accordance with the above RT-qPCR and IF analyses. Downregulated genes in HC-CD56^neg^ *vs*. HC-CD56^pos^ cells, that identified genes which expression was reduced at the time of the loss of myogenicity in normal cells, were related to chromatin organization, protein-DNA complex and DNA and chromatin binding (Fig. 3A, red label in the left box). When comparing DMD-CD56^pos^ *vs*. HC-CD56^pos^, to identify genes that were differentially expressed in DMD *vs.* HC myogenic cells, we observed that the down-regulated genes were also related to chromatin organization, regulation of DNA binding, and chromatin (Fig. 3A, red label in the right box). Scrutinizing those two lists of down-regulated genes, 11 common genes were found (Fig. 3B). RT-qPCR experiments run on HC-CD56^pos^ and DMD-CD56^pos^ cells confirmed the differential expression of 4 of them. They included: *CBX3*, encoding for the heterochromatin protein HP1γ, *H2AZ2* and *H3F3B* (encoding for H3.3), two histone variants, and *SMC3*, a subunit of the chromatin cohesion complex (Fig. 3C).

**Fig. 3.**
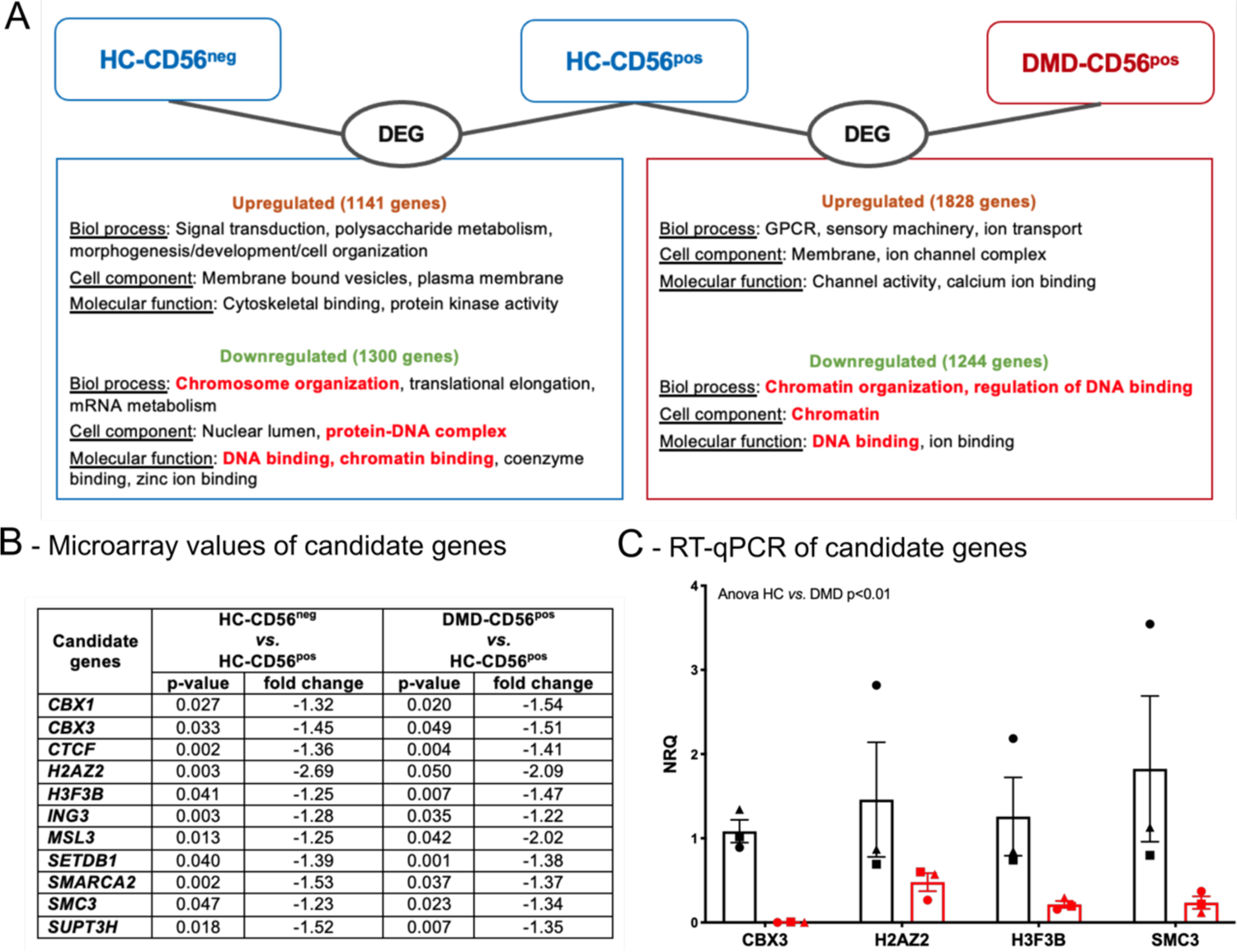
Transcriptomic analysis of CD56^pos^ and CD56^neg^ cells from HC- and DMD-MuSC cultures. **(A)** Gene ontology *(DAVID software)* of differentially expressed genes (DEG) after microarray analysis of CD56^pos^ and CD56^neg^ cell*s* issues from healthy control (HC)- and Duchenne (DMD) CD56^pos^ MuSC cultures as depicted in Fig. 2A. **(B)** Microarray fold change of the 11 genes found down expressed in both HC-CD56^neg^ *vs*. HC-CD56^pos^ and DMD-CD56^pos^ *vs*. HC-CD56^pos^ cells. **(C)** Normalized Relative Quantity (NRQ) expression by RT-qPCR of *CBX3*, *H2AZ2*, *H3F3B* and *SMC3* genes in HC- and DMD-CD56^pos^ cells. Results from 3 HC and 3 DMD samples. Each shape symbol represents the same culture.

These results indicate that the expression of 4 epigenetic regulators was decreased at the time of the myogenic loss in healthy MuSCs, and was already low in myogenic DMD-MuSCs, as compared with HC MuSCs.

### Lentiviral expression of *CBX3*, *H2AZ2*, *H3F3B* and *SMC3* rescues the myogenic potential of DMD-CD56^pos^ cells

In order to reexpress the 4 genes *CBX3*, *H2AZ2*, *H3F3B* and *SMC3* in DMD-MuSCs, a protocol of exogenous expression of the 4 genes with a hemagglutinin (HA) reporter sequence using lentiviral transduction was designed (Fig. 4A). The EF1α promoter was selected to ensure a stable long-term expression of the transduced genes (Massenet et al. 2020). After transduction, cells were selected with puromycin and were cultured for 5 to 10 population doublings (*i.e.* the time for having around 50% loss of CD56 expression in DMD-MuSCs) before analysis. Immunostaining for HA confirmed the expression of the lentiviruses in MuSCs in all conditions but the control (Fig. S4A). The expression of the 4 genes was evaluated by RT-qPCR and showed a robust increase of their expression, albeit some high variations were observed between some samples (Fig. S4B).The expression of the *CBX3*, *H2AZ2*, *H3F3B* and *SMC3* or of the 4 genes together in DMD-MuSCs was associated with an increase of the number of cells expressing CD56 after several weeks in culture as compared with cells transfected with an empty virus (Fig. 4B). The mean increase of CD56 expression ranged from 20 to 31%. Additionally, the simultaneous transduction of DMD-CD56^pos^ cells with the 4 genes together induced a higher increase in the number of CD56^pos^ cells, of about 60%, as compared with the control (Fig. 4B). Then, the capacity of the transduced cells to implement myogenesis was evaluated (because of the scarcity of the material, only differentiation assay was carried out, fusion assay requiring too many cells). Consistent with their increased CD56 expression, transduced DMD-CD56^pos^ increased their expression of myogenin (Fig. S4C). These results show that the re-expression of *CBX3*, *H2AZ2*, *H3F3B* and *SMC3* genes in DMD-MuSCs counteracted the loss of CD56 expression and maintained their myogenic identity *in vitro*.

**Fig. 4.**
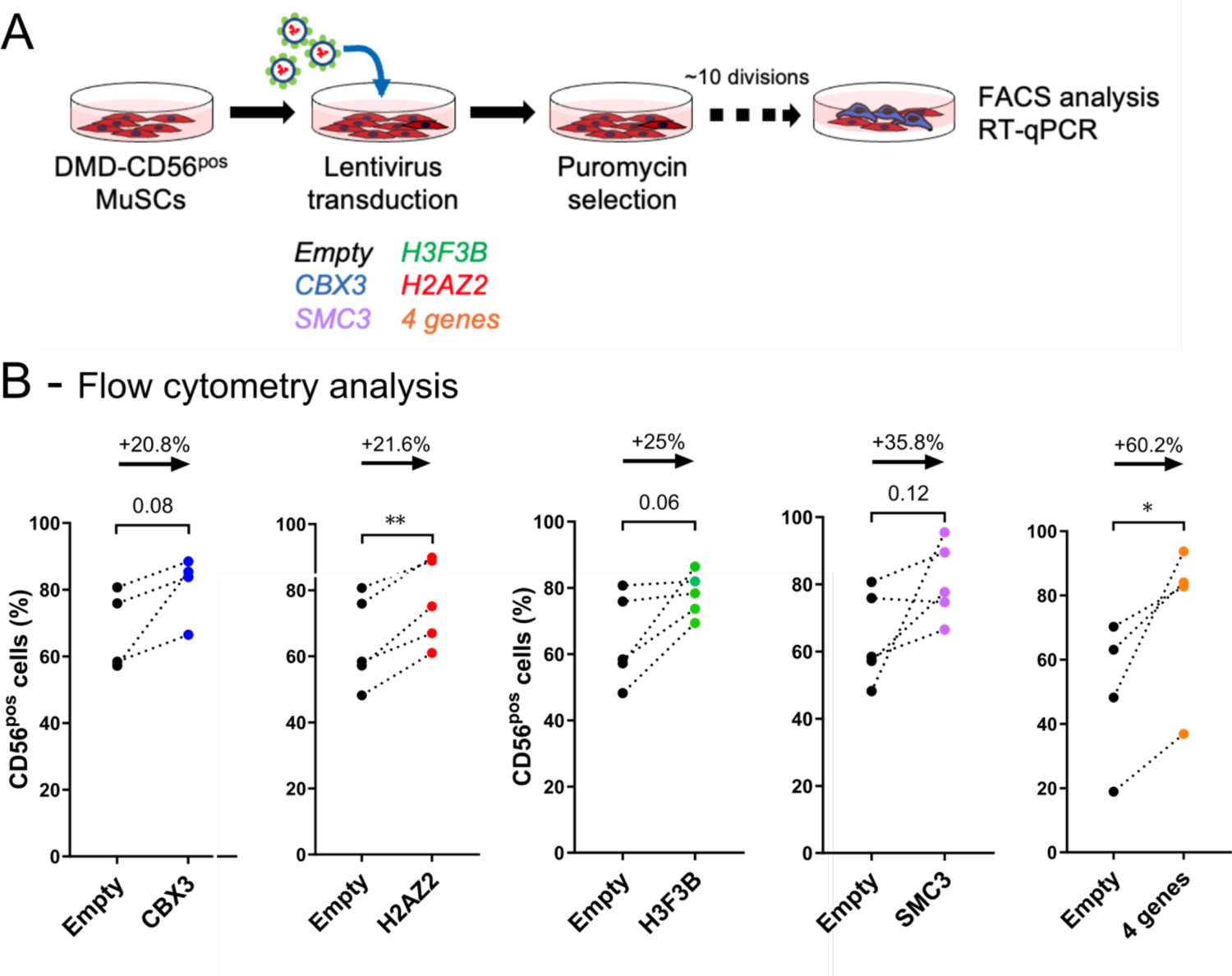
Transduction of DMD-CD56^pos^ cells with *CBX3*, *H2AZ2*, *H3F3B* and *SMC3* coding lentiviruses. **(A)** Experimental procedure for DMD-CD56^pos^ cell transduction with lentiviruses. Cells were cultured in growth medium. **(B)** Flow cytometry quantification of the number of CD56^pos^ cells in each condition. Results are from 4 to 5 DMD samples. Results are means ± SEM of 3 DMD samples (each shape symbol is used for cells issued from the same initial culture). *p<0.05, **p<0.001 using paired t-test.

### ATACseq identifies a target of CBX3, H2AZ2, H3.3 and SMC3 in MuSCs

To identify potential targets of CBX3, H2A.Z2, H3.3 and SMC3 in MuSCs, ATAC sequencing was performed before and after lentiviral transduction. Five conditions were analyzed (Fig. 5A): (1) pure DMD-CD56^pos^ cells harvested before the infection (Day 0); (2) empty-CD56^pos^ and (3) empty-CD56^neg^ cells resulting from DMD-CD56^pos^ cells transfected with empty lentivirus and cultured for about 10 population doublings; (4) 4V-CD56^pos^ and (5) 4V-CD56^neg^ cells resulting from DMD-CD56^pos^ cells transfected with *CBX3*, *H2AZ2*, *H3F3B* and *SMC3* lentiviruses and cultured for 10 population doublings.

**Fig. 5.**
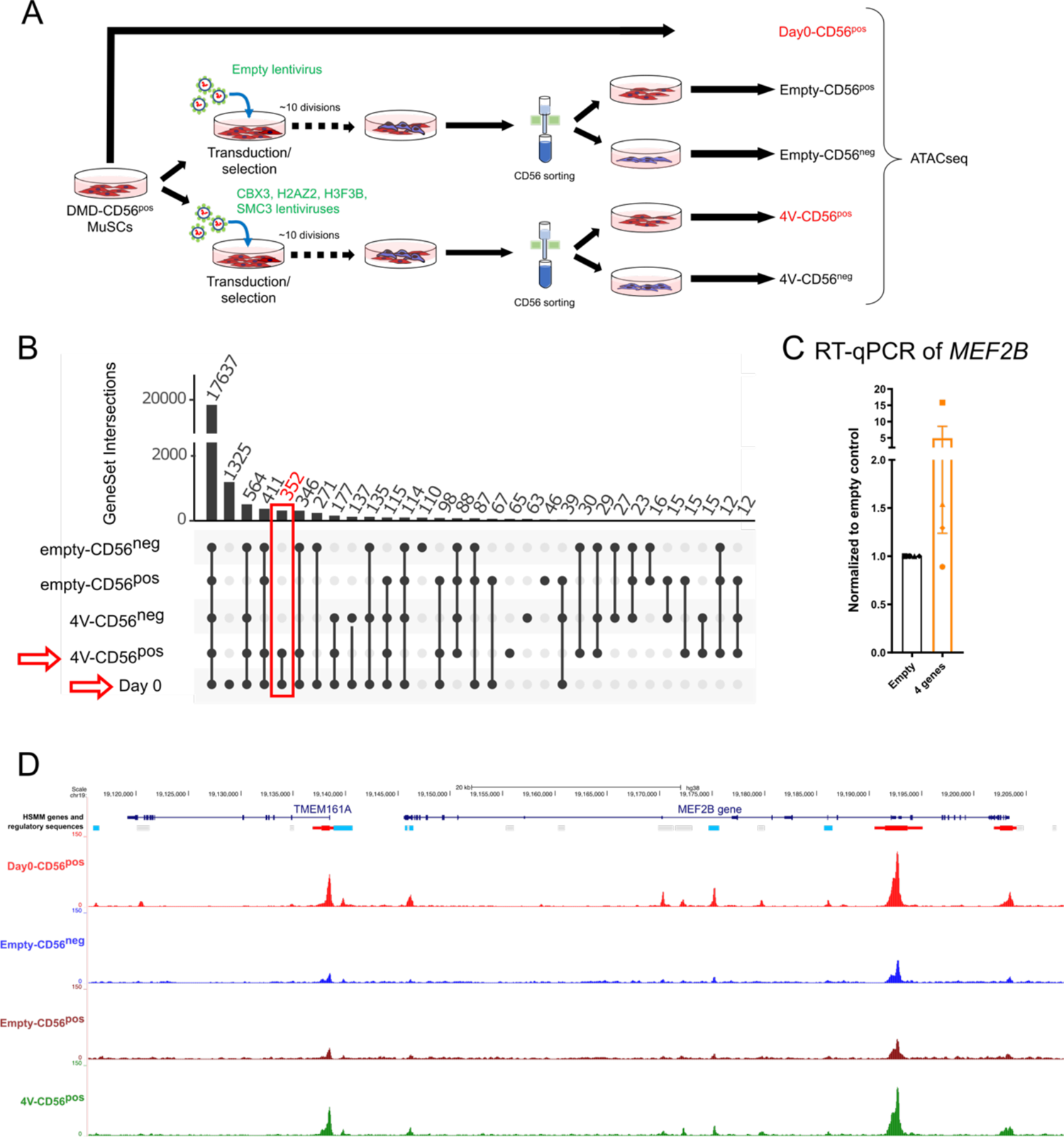
Identification of CBX3, H2AZ2, H3.3 and SMC3 target genes using ATAC-seq. **(A)** Experimental design of ATAC-seq analysis of DMD-CD56^pos^ and DMD-CD56^neg^ cells initially non-transduced (Day0-CD56^pos^) or transduced with either empty (empty-CD56^pos/neg^) or *CBX3*, *H2AZ2*, *H3F3B* and *SMC3* lentiviruses together (4V-CD56^pos/neg^). **(B)** UpSet plot of the comparison between the samples to identify genes differentially expressed in cells transduced with the 4 epigenetic regulators and non-transduced (red arrows) *versus* the 3 other conditions (red rectangle). **(C)** Expression by RT-qPCR of the *MEF2B* gene in DMD-CD56^pos^ cells transduced with *CBX3*, *H2AZ2*, *H3F3B* and/or *SMC3* lentiviruses. **(D)** Screenshot of *MEF2B* locus. From top to bottom: chromosome scale, gene and regulatory sequences, ATACseq tracks in CD56^pos^ DMD-MuSCs initially non-transduced, DMD-CD56^pos^ and DMD-CD56^neg^ cells from CD56^pos^ DMD-MuSCs initially transduced with an empty vector, and DMD-CD56^pos^ cells from CD56^pos^ DMD-MuSCs initially transduced with the 4 epigenetic regulators (see Fig. 5A). Results are means ± SEM of 4 samples.

Each ATAC seq peaks obtained from the five conditions were associated to the unique closest gene to obtain a list of genes. Next, gene list comparison was performed to obtain a list of genes that were present in both Day 0 and 4V-CD56^pos^ samples but that were absent in empty-CD56^pos^, empty-CD56^neg^ and 4V-CD56^neg^ samples (red arrows in Fig. 5B). These 352 genes supposedly presented DNA sequences with open chromatin in both DMD-CD56^pos^ cells before the loss of CD56 and in cultured 4V-CD56^pos^ cells when accessibility was maintained thanks to the exogenous expression of CBX3, H2AZ2, H3.3 and SMC3 proteins. This list was cross-analyzed with the list of downregulated genes between DMD-CD56^neg^ and HC-CD56^pos^ cells identified by transcriptomics (Table S2).

This resulted in a list of 78 genes defined as down-regulated in DMD-CD56^pos^ cells and potential targets of CBX3, H2A.Z2, H3.3 and SMC3 in the ATAC seq analysis (Fig. S5A). Among them, only 34 genes presented an ATACseq peak at a regulatory sequence (Fig. S5A). This list of 34 genes was refined by cross-analysis with ENCODE polyA-RNAseq from normal human primary myoblasts (ENCSR000CWN) and allowed to retrieve 8 candidates: *ABHD14A*, *APBA3*, *EXOSC10*, *IF172*, *MEF2B*, *MIOS*, *NMP3* and *TNKS* (Fig. S5A). The expression of the 8 genes was evaluated by RT-qPCR in DMD-MuSCs after lentiviral transduction of *CBX3*, *H2AZ2*, *H3F3B*, *SMC3* and of the 4 genes together (Fig. S5B). Only *MEF2B* showed a robust increased expression in all conditions (Fig. 5D and S5C), which was confirmed by an increase in chromatin accessibility at the *MEF2B* locus observed by an higher ATACseq peak intensity after the lentiviral transduction of all 4 genes together in DMD-MuSCs. (Fig. 5D).

### MEF2B is a target of H2AZ2 and H3.3 in MuSCs and is required for the maintenance of the myogenic identity

To confirm that MEF2B was a target of CBX3, H2AZ2, H3.3 and SMC3, Cut&Tag technology was used to evaluate protein binding at the locus after lentiviral transduction of DMD-CD56^pos^ cells with lentiviruses expressing the 4 genes (Fig. 6A). Results showed the presence of the H2AZ2 histone variant at the promoter of the *MEF2B* gene (Fig. 6B) and the presence of both H2AZ2 and H3.3 histone variants at the enhancer of the *MEF2B* gene (Fig. 6B).

**Fig. 6.**
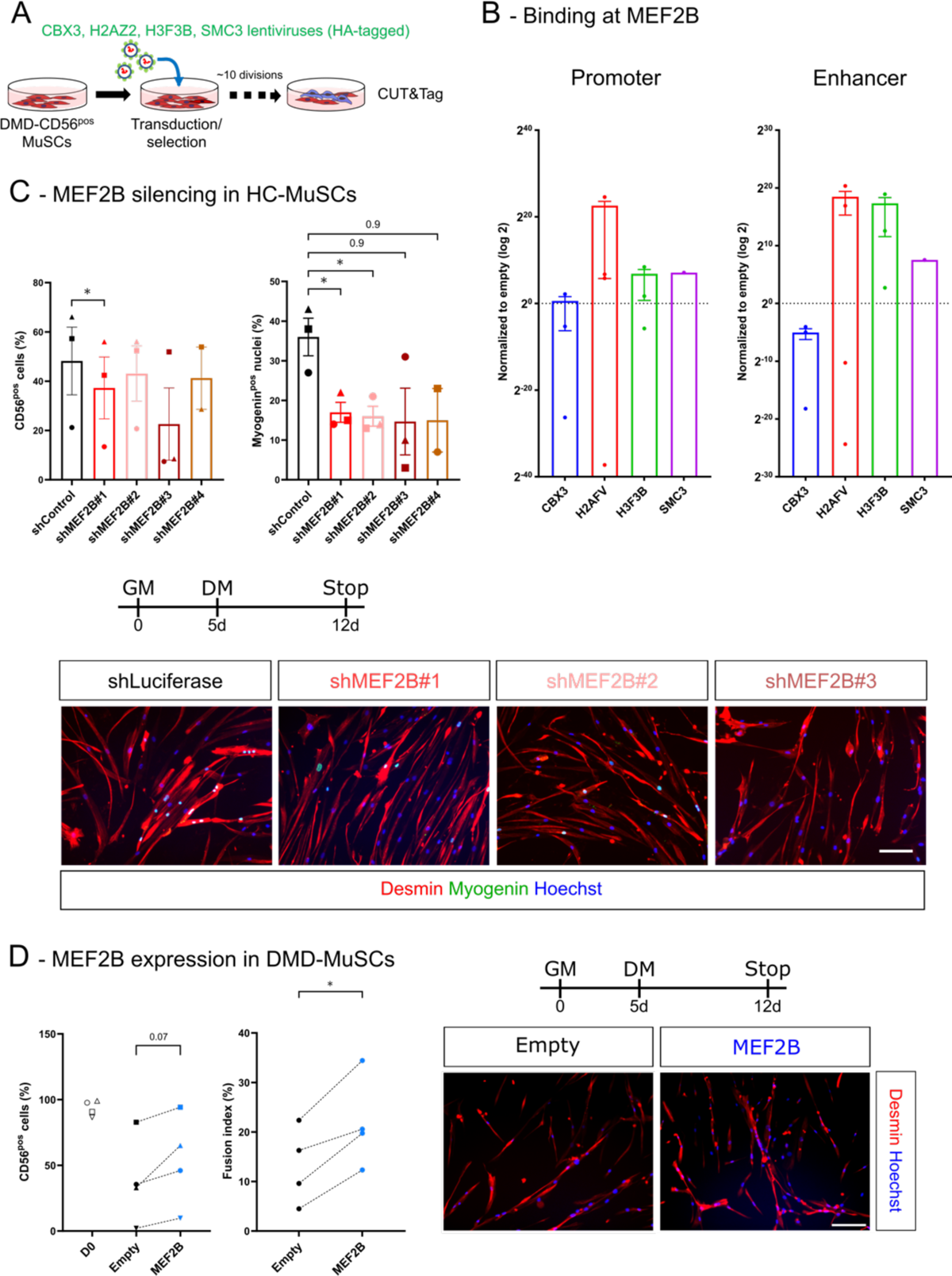
MEF2B and the maintenance of MuSC myogenicity. **(A)** Experimental procedure for DMD-CD56^pos^ cell transduction with empty or HA-tagged *CBX3*, *H2AZ2*, *H3F3B* or *SMC3* coding lentiviruses before the Cut&Tag analysis. Cells were cultured in growth medium. **(B)** Normalized Relative Quantity expression of MEF2B promoter and enhancer sequences by RT-qPCR after Cut&Tag library preparation with DMD-CD56^pos^ cells transduced as depicted in (A) using anti-HA antibodies to target the 4 epigenetic regulators. Dotted lines correspond to the expression after using control IgGs. **(C)** Loss of function experiments where HC-CD56^pos^ were transduced with 4 different shRNA*MEF2B* lentiviruses (and shLuciferase as a control) and were analyzed for their CD56 expression by flow cytometry and for their myogenic capacity, assessed by myogenin expression (green) when cultured in differentiation medium (DM) (desmin is red, nuclei are blue [Hoechst]). Data are means ± SEM of 2 to 3 experiments. **(D)** Gain of function experiments where DMD-CD56^pos^ cells were transduced with a lentivirus encoding for *MEF2B* and were analyzed for their CD56 expression by flow cytometry and for their myogenic capacity, assessed by their fusion index when cultured in differentiation medium (desmin is red, nuclei are blue [Hoechst]). Data are shown for 4 DMD donors. *p<0.05,using paired t-test. Bars = 100 μm.

To further explore the function of MEF2B in the maintenance of the myogenic identity of CD56^pos^ cells, we performed loss and gain of function experiments. shRNA lentivirus against *MEF2B* was transduced in HC-CD56^pos^ cells and the percentage of CD56^pos^ cells was evaluated after 10 population doubling following the puromycin selection (efficacy of the shRNAs on MEF2B expression is shown in Fig. S5C). *MEF2B* knockdown using 4 different shRNAs induced a decrease of the percentage of CD56^pos^ cells as compared with the control shRNA (Fig. 6C). Functionally, the myogenesis assay showed that the percentage of myogenin^pos^ cells was strongly reduced in sh*MEF2B* treated HC-MuSCs, by 52-58% (Fig. 6C). Gain of function experiments included the transduction of DMD-CD56^pos^ cells with a *MEF2B* lentivirus as described above for the 4 epigenetic regulators. The increase of *MEF2B* expression by transduced DMD-MuSCs (Fig. S5C) was associated with an increase of the number of cells that expressed CD56 (+142%) (Fig. 6D). Functionally, the DMD-MuSCs expressing *MEF2B* showed an increased capacity to differentiate and to form myotubes in vitro (+165%) (Fig. 6D). These results show that in DMD cells, alterations in the chromatin organization prevented the expression of *MEF2B* and that MEF2B was involved in the maintenance of the myogenic identity of MuSCs.

## Discussion

In the present study, we have examined how the DMD pathology affects the lineage fidelity of MuSCs. We found that human MuSCs isolated from DMD muscle continue to show myogenic properties that allow them to self-renew and differentiate to form new myofibers. Nevertheless, the overall MuSC population showed a rapid decline in their myogenicity where they drifted towards a fibroblast-like cell identity. This change in cell identity was observed at the clonal level, and resulted from the altered expression of epigenetic enzymes required to maintain the myogenic cell fate. Amongst the epigenetic changes, a closing of chromatin at the gene encoding the transcription factor MEF2B caused a down-regulation of its expression, and a loss of the myogenic fate. Thus, our work identified MEF2B as a key mediator of the myogenic fate in human MuSCs.

The continued expansion of both healthy and DMD human MuSCs resulted in a loss of the myogenic identity in favor of a more fibroblast-like identity. Differential analysis of gene expression identified 4 important epigenetic factors (*CBX3*, *H2A.Z2*, *H3.3*, and *SMC3*) as being down-regulated during the same time frame, suggesting that an epigenetic drift may be at the heart of this loss of myogenicity. The down-regulation of each of these 4 factors is likely to contribute to the loss of myogenic gene expression. The SMC3 protein is a subunit of the cohesion complex, a key mediator of DNA looping that allows the communication between transcriptional enhancers and promoters to facilitate transcription (Hansen et al. 2018; Dall’Agnese et al. 2019). In the absence of SMC3, a loss of Topological Associated Domains (TADs) in the DMD-MuSCs would prevent the muscle-specific enhancers from communicating with promoters, and would result in a reduction of muscle gene expression. Similarly, CBX3 is required to maintain high levels of muscle gene expression as the euchromatin-associated protein helps facilitate transcriptional elongation by promoting RNA Polymerase II pause-release and recruitment of the FACT complex required for removal of nucleosomes that impede polymerase progression (Yahi et al. 2008; Kim et al. 2011; Zhong et al. 2019; Schoelz and Riddle 2022). Finally, the histone variants H2A.Z2 and H3.3 proteins promote a transcriptionally permissive chromatin state by establishing a less stable nucleosome that is easily displaced by chromatin remodeling factors to establish open chromatin (Santisteban et al. 2000; Jin et al. 2009; Vardabasso et al. 2015). Interestingly, H3.3 been shown to contribute to cellular memory where methylation of the histone variant at the K4 position allowed reprogrammed cells to “remember” their myogenic cell identity (Ng and Gurdon 2008; Harada et al. 2012). H3.3 plays a dual role in maintaining cell memory by both promoting the muscle gene regulatory program and suppressing genes of alternate lineages. Indeed, when H3.3 is not incorporated into the genome due to loss of its chaperon protein HIRA, MuSCs begin to express non-muscle genes that are normally restricted to alternate lineages in (Esteves de Lima et al. 2021). Thus, the loss of myogenicity observed due to the continued expansion of DMD-MuSCs is likely due to epigenetic drift where myogenic genes become repressed while genes of the fibroblast lineage become expressed.

Our study has identified MEF2B as a key myogenic gene turned off during DMD-induced epigenetic drift. While MEF2 family of proteins have been widely studied in muscle, the MEF2B protein has largely been overlooked as a contributor to the myogenic fate. Among MEF2 family members, MEF2B presents a unique protein structure and doesn’t bind the MEF2 consensus DNA motif because of the presence of its C-terminal domain (Molkentin et al. 1996). In addition, MEF2B expression in the heart and muscle of adult mice is low compared to the other MEF2 proteins while MEF2B is highly express in human adult muscles (Morisaki et al. 1997). Despite its limited role in murine myogenesis, we find that MEF2B is essential to maintaining the myogenic cell fate. Indeed, exogenous expression of MEF2B in DMD-MuSCs reduced the myogenic loss of identity while depletion of MEF2B in MuSCs increased the rate at which the myogenic identity was lost. Interestingly, previous studies have hinted at a role for MEF2B in myogenesis. Co-expression of MEF2B with either Pax3 and Pixt1 or Pax7 and Pixt1 can promote differentiation of embryonic fibroblast into MuSCs (Ito et al. 2017). Moreover, MEF2B expression associated with Pax7 and MyoD can also promote transdifferentiation of adult fibroblasts in MuSCs (Ito et al. 2017). These findings suggest that MEF2B could be a gatekeeper of the myogenic lineage during adult myogenesis.

It is unclear whether the accelerated epigenetic drift observed in DMD-MuSCs is due to the loss of dystrophin itself, or due to downstream signaling initiated by the loss of dystrophin. In the absence of dystrophin in myofibers leads to miss localization of the Dystrophin Associated Protein Complex (DAPC) and the loss of Nitric Oxide Synthase (NOS) signaling (Brenman et al. 1995; Lai et al. 2009). NOS signaling is necessary to regulate chromatin accessibility with the regulation of Histone Deacetylases (HDACs) in other different cell types (Nott et al. 2008; Vasudevan et al. 2016; Mengel et al. 2017). In DMD myofibers, the absence of NOS signaling leads to aberrant activation of HDAC2 and global changes of histone acetylation across the genome (Illi et al. 2008; Colussi et al. 2009). The DAPC also regulates MuSCs polarity and asymmetric division at the epigenetic level (Dumont et al. 2015b; Chang et al. 2018). Moreover, in addition to the aberrant histone modification caused by the absence of dystrophin, it also leads to a deregulated ncRNA and genomic instability (Cacchiarelli et al. 2010; Schmidt et al. 2011; Iyer et al. 2019). While we observe that DMD results in an epigenetic silencing of MEF2B expression it remains unclear why the chromatin associated with this locus becomes transcriptionally repressive. However, it will be important to understand how this epigenetic drift occurs in DMD since many of the current therapy trials look to restore Dystrophin expression, and have not examined restoration of downstream epigenetic pathways.

An important finding of our studies is that DMD-MuSCs initially possess normal myogenic potential, but that this potential is lost over time due to epigenetic drift. While the properties of MuSCs have been extensively investigated, controversy in the field has persisted about the myogenic properties of MuSCs. Indeed, the study of DMD were hindered by two major hurdles: i) investigation in mouse, using the mdx model, which poorly recapitulates DMD features and ii) investigation using human cells, that are no longer in their in vivo environment, with a risk of developing culture-driven artifacts. In its infancy, culture of human MuSCs was further limited by the presence of non-myogenic cells in the isolated cell population, that were bulk cultured. Nevertheless, two types of cells were observed in those cultures, including myogenic cells, and cells that do not fuse and exhibit a fibroblastic phenotype, the number of which is increased in DMD samples as compared with normal muscle (Yasin et al. 1979; Blau et al. 1983b; Delaporte et al. 1984). Clonal cultures confirmed the presence of non-fusing low creatine kinase activity cells along with myogenic cells, capable of fusion (Webster and Blau 1990; Meola et al. 1991). From the initial muscle samples, less cloneable MuSCs were obtained in DMD versus normal muscle (Webster and Blau 1990; Meola et al. 1991) indicative of an enrichment of the diseased muscle by fibroblastic cells. Careful examination of the cell phenotype of unsorted cells, then culture of cells that were purified according to the CD56 expression showed that myogenic cells implement of normal myogenic process in DMD as compared with normal muscle, including proliferation, differentiation and fusion (Blau et al. 1983a; Zanotti et al. 2007). In the present study, we also demonstrated that MuSCs that express CD56, and thus be considered as myogenic (Illa et al. 1992), show unaltered myogenic properties. Similar results were observed in the large animal model GRMD dog (Berg et al. 2011). Thus, as long as they are isolated as myogenic cells (expressing CD56), DMD-derived MuSCs have retained their full myogenic capacities.

Using purified CD56^pos^ MuSCs cultures, we observed a progressive reduction in the proportion of myogenic cells with time. The loss of myogenicity has been repeatedly observed by other labs in cultures of human MuSCs isolated from normal (healthy control, HC) muscle, with very high variations between donors, and independently of their age and sex (Agley et al. 2013; Alsharidah et al. 2013; Francis et al. 2022). However, our results showed that the decrease in the proportion of myogenic CD56^pos^ cells was twice as fast in DMD cultures as compared with HC cultures. No difference was observed in the growth rate or apoptosis between CD56^pos^ and CD56^neg^ cell populations of both DMD and HC cell cultures, suggesting a transition of myogenic cells into fibroblastic cells with time. To validate this transition, we made clonal cultures of single CD56^pos^ cells that expressed Myf5 and found that these myogenic cells lose their myogenic nature and acquired the feature of fibroblasts by TCF7L2 expression while losing that of CD56. A few cells are double positive, suggesting these cells are in transit between the two statuses. Such a transition was observed in normal MuSCs (Agley et al. 2013; Francis et al. 2022) where CD56^neg^ cells are capable of adipogenic differentiation while CD56^pos^ are not (Agley et al. 2013; Francis et al. 2022). After the same period of time, DMD-MuSC derived clones exhibit 3-fold more CD56^neg^ cells than HC-MuSC-derived cells, confirming that the transition occurs much faster in DMD cultures. This transition is associated with a general downregulation of the expression of several myogenic genes (*PAX7*, *ACTA1 MYF5*, *MYOD*) with a concomitant increased expression of genes linked to a fibroblastic phenotype (*COL1A1*, *CTGF*, *LOX*, *SPP1*). Such increase of gene associated with matrix deposition and matrix remodeling was previously reported in CD56^pos^ human MuSCs isolated from DMD muscle (Zanotti et al. 2007). Whether this transition occurs in human in vivo is impossible to address. However, the presence of MuSCs expressing a canonical marker of fibrogenic cells (Pax7^pos^; PDGFRα^pos^) was reported in human DMD muscle (Pessina et al. 2015). Moreover, in the mdx mouse, such a fibrogenic plasticity has been observed. A part of MuSCs lose their myogenic nature to acquire a fibrogenic identity, driven by a Wnt-TGFβ axis (Biressi et al. 2014; Pessina et al. 2015). Thus, there is evidence that indicates that MuSCs may undergo epigenetic drift in vivo towards a fibroblast-like identity.

## Materials and Methods

### Patients and primary cultures

Biopsies were obtained from *deltoideus medialis* of 21 genetically characterized DMD patients. Fourteen patients undergoing orthopedic surgery (intercostal muscle) or for which the deltoid biopsy showed no signs of neuromuscular diseases and for whom the diagnosis workup was normal were used as age-matched controls. Cells were recovered from the hospital cell bank (protocol registered at the Ministère de la Recherche and Cochin Hospital Cell Bank, Paris, agreement n°DC-2009-944). From the muscle biopsy to the delivery by the cell bank, muscle cells were expanded for about 7-10 days before magnetic cell sorting based on CD56 expression was performed (Fig. S1A). Purity of the cells was evaluated after flow cytometry evaluation of CD56 expression (see below) and by immunofluorescence (IF) after 6h of culture on glass coverslips. After fixation (paraformaldehyde [PFA] 4%) and permeabilization (triton X-100 0.5%), cells were incubated with anti-CD56 mouse antibodies (1:10, Coulter Clone #PN4235479-G) and anti-Pax7 rabbit antibodies (1:200, Abcam #ab92317) overnight at 4°C, that were revealed by Cy3-coupled anti-mouse IgGs (1:200, Jackson ImmunoResearch #715-165-150) and Cy5-coupled anti-rabbit IgGs (1:200, Jackson ImmunoResearch #711-165-152) for 45 min at 37°C. Nuclei were labeled with Hoechst (Sigma #B2261) and mounting was done in Fluoromount (Sigma #F4680).

### Culture of primary human MuSCs

Primary cells were cultured in growth medium, which includes Skeletal Muscle Cell Growth Medium (Promocell #C23260) containing skeletal muscle supplemental mix (Promocell #C39365), 10% of Fetal Bovine Serum (FBS) (Abcys #S1810-500), 100 U/ml penicillin and 100 µg/ml streptomycin (Gibco #15140).

### Evaluation of CD56 expression by flow cytometry

Trypsinized or MACS sorted cells were incubated at 4°C for 20 min with APC-conjugated anti-CD56 antibodies (1:40, BD Pharmingen #555518) or isotype control (1:40, BD Pharmingen #555751) and further analyzed using a FACSCanto II flow cytometer (BD Biosciences).

### Immuno-magnetic cell sorting

Cells were trypsinized, centrifuged and resuspended in 170 µl of magnetic-activated cell sorting (MACS) buffer (Phosphate-Buffered Saline solution (PBS) containing 5% Bovine Serum Albumin (BSA, Sigma #A9647), 1 mol/l of Ethylenediaminetetraacetic Acid Disodium (EDTA, Sigma #ED2SS). Thirty µl of superparamagnetic microbeads conjugated to a CD56 primary antibody (Miltenyi #130-050-401) was mixed with the cell suspension and incubated for 30 min at 4 °C. After a wash with MACS buffer, the cell suspension was passed through a 30 µm filter (Miltenyi #130-041-407) and dripped into a LC column (Miltenyi #130-042-202) held in a MiniMACS magnetic separation unit (Miltenyi). Eluted cells were recovered as CD56^neg^ cells. Column was then removed from the magnetic separation unit, and flushed with MACS buffer to recover CD56^pos^ cells.

### Proliferation assay

CD56^pos^ cells were seeded at 3,000 cells per cm^2^ in 4 well Permanox Nunc Lab-Tek chambers (ThermoFisher #177437) and cultured in growth medium. Two days later, EdU (Click-iT EdU Alexa Fluor 488 Imaging Kit, ThermoFisher #C10337) was added at 1 µg/ml and cells were further incubated for 8 h and then processed following the manufacturer’s instructions. Nuclei were labeled with Hoechst and mounting was done in Fluoromount. The percentage of proliferating cells was calculated as the number of Alexa fluor-488 positive nuclei over the total number of nuclei.

### Myogenesis assay – differentiation

Freshly CD56^pos^ sorted cells were seeded at 1,000 cells per cm^2^ in 4 well Permanox Nunc Lab-Tek chambers in growth medium. Six h later, medium was replaced by differentiation medium (Skeletal Muscle Basal Medium containing 10 µg/ml human insulin [Sigma #I2643], 100 U/ml penicillin and 100 µg/ml streptomycin) and cells were cultured for 5 days. Long term cultured transduced cells were seeded at 10,000 cells per cm^2^ in 8 well Permanox Nunc Lab-Tek chambers (ThermoFisher #177402) and grown for 5 days in growth medium. Then, differentiation medium was added and cells were further incubated 7 days. IF was performed as described above using anti-myogenin mouse antibodies (1:50, BD Biosciences #556358) and anti-desmin rabbit antibodies (1:200, Abcam #ab32362) revealed by Cy3-coupled anti-mouse IgGs and Cy5-coupled anti-rabbit IgGs. The percentage of differentiated cells was calculated as the number of cells positive for myogenin nuclei over the total number of cells.

### Myogenesis assay – fusion

Freshly CD56^pos^ cells were plated at 500 cells per cm^2^ in 175 cm^2^ Nunc Flask (ThermoFisher #156502) in growth medium. Six h later, medium was replaced by differentiation medium and cells were cultured for 5 days. Then, these differentiated cells were trypsinized and seeded at 50,000 cells per cm^2^ in 8 well permanox Nunc Lab-Tek chambers in growth medium. Six h later, growth medium was replaced by differentiation medium and cells were cultured for 3 days. Long term cultured transduced cells were seeded at 10,000 cells per cm^2^ in 8 well permanox Nunc Lab-Tek chambers. After 5 days of culture in growth medium, differentiation medium was added and cells were cultured 7 days more. IF was performed as described above using anti-desmin rabbit antibodies revealed by Cy5-coupled anti-rabbit IgGs. The fusion index was calculated as the number of nuclei in cells presenting two or more nuclei over the total number of nuclei.

### TUNEL assay

Cells were seeded at 3,000 cells per cm^2^ in 12-well plates containing glass coverslips and were incubated for 24 h. After PFA fixation and Triton permeabilization, cells were stained for DNA strand break using Click-iT^TM^ Plus TUNEL Assay (ThermoFisher #C10617) following manufacturer’s protocol before mounting in Fluoromount.

### Plasmid construction

The pLenti-GIII-EF1α-linker-Flag-Hemagglutinin(HA) plasmid was built by insertion of a linker sequence containing cutting sites in the pLenti-EF1α-SetD2-HA plasmid (ABMgood #435220610695), digested at cutting sites of EcoRV, with CloneEZ PCR Cloning Kit (GenScript #L00339). The pLenti-GIII-EF1α-CBX3-Flag-HA, pLenti-GIII-EF1α-H2AZ2-Flag-HA, pLenti-GIII-EF1α-H3F3B-Flag-HA and pLenti-GIII-EF1α-SMC3-Flag-HA plasmids were built by insertion of a PCR amplified CBX3, H2AZ2, H3F3B or SMC3 gene cDNA in the pLenti-EF1α-linker-Flag-HA plasmid, digested at cutting sites BamHI and KpnI for H2AZ2, NheI and XbaI for H3F3B, of EcoRV, using the CloneEZ PCR Cloning Kit. The PCR amplification of CBX3, H2AZ2, H3F3B or SMC3 was performed with Phusion High-Fidelity DNA Polymerase (NEB #M0530S) on human MuSC cDNA. The cDNAs were built by reverse transcription of proliferating MuSC mRNAs using M-MuLV Reverse Transcriptase (NEB #M0253L). mRNAs were extracted using NucleoSpin RNA Plus XS kit (Macherey-Nagel #740990.50). MEF2B was cloned in pLenti-GIII-EF1α-HA plasmid by abmgood using NheI and BamHI cutting sites (ABMgood, #2832406). The pLV-Myf5Promoter-GFP plasmid was built by insertion of two sequences in a pLKO.1 plasmid backbone (Addgene #10878). The Myf5-promoter sequence was added at NdeI and KpeI cutting sites and the CRE-GFP sequence was added at SpeI and XhoI restriction sites using a T4 DNA ligase (NEB #M0202L). Linker sequence and primers used for PCR amplification are listed in Table S3. The MEF2B shRNA sequences inserted in a pLKO-1 puro plasmid are listed in Table S4 (Sigma, #SHCLNG).

### Lentiviral production

Lentiviral production was carried out by CaCl2 transfection of HEK293T cells (1×10^6^ cells) using 19.9 µg of constructed lentiviral vector or a pLenti-GIII-EF1α empty (ABMgood #LV588), 5.93 µg of MD2.G plasmid (Addgene #12259) and 14.88 µg of psPax2 plasmid (Addgene #12260) during 15 h. Supernatant containing lentivirus was collected 24 and 48 h after the end of transfection, filtered and concentrated using sucrose buffer and ultracentrifugation at 120,000 g for 2 h at 4°C. Lentiviral titration was estimated by transfection of healthy MuSCs with concentrated lentivirus in growth medium supplemented by 6 µg/ml of polybren (Sigma #107689). One day after transfection, cells were selected in growth medium supplemented with 1 µg/ml puromycin (Sigma #P8833). Multiplicity of Infection (MOI) was calculated by counting the remaining cells 3 days after the start of selection.

### MuSC lentiviral transduction and selection

Cells were seeded at 3,000 cells per cm^2^ in 6-well plates in growth medium and 6 h later, medium was replaced by growth medium supplemented with 6 µg/ml polybren and lentivirus to a final MOI of 1. After 36 h, medium was replaced by selection medium (growth medium containing 1 µg/ml puromycin). After 3 days of puromycin selection, MuSCs were cultured in growth medium for 5-10 divisions for further analysis. Transduction efficacy was validated by IF for HA using anti-HA antibodies (1:200, Roche, 11583816001) according to the IF protocol described above.

### Clonal cell culture

After LV-Myf5Promoter-GFP lentivirus transduction, cells were labeled with APC-conjugated anti-CD56 antibodies. Cells were sorted using a BD FACSAria II for GFP^pos^/CD56^pos^ cells, that were clonally seeded at one cell per well in 96-well plates coated with Matrigel (Corning Life Sciences) (Matrigel:conditioned growth medium [1v:99v]). Conditioned growth medium was recovered every 24h from healthy immortalized MuSCs (Massenet et al. 2020). Clones were cultured in conditioned medium until they reach the density of 3,000 cells per cm^2^ and were thereafter cultured in growth medium for 4-6 weeks. Cells were proceeded for IF as described above using anti-CD56 mouse antibodies (1:10, Coulter #PN4235479-G) and anti-TCF7L2 rabbit antibodies (1:100, Cell Signaling Technology #C48H11), revealed by Cy3-coupled anti-mouse IgGs and Cy5-coupled anti-rabbit IgGs.

### Quantitative RT-PCR

Total RNAs were extracted using NucleoSpin® RNA Plus XS kit (Macherey-Nagel #740990.50). The quality of RNA was checked using Nanodrop. One microgram of total RNA was reverse-transcripted using Superscript II Reverse Transcriptase (ThermoFisher #18064022) and diluted 5 times. Each sample was tested in triplicate. Quantitative RT-PCR was performed using CFX96 Real-Time PCR Detection System (Bio Rad). The 10 µl final volume of reactive mixture contained 2 µl of diluted cDNA, 0.5 µl of primer mixture (Table S5), 2.5 µl of water and 5 µl of LightCycler 480 SYBR Green I Master Kit (LifeSciences #04707518001). After initial denaturation of 2 min, the amplification was performed for 45 cycles of 95°C for 10 sec, 60°C for 5 sec and 72°C for 10 sec. The calculation of normalized relative quantity (NRQ) was performed using *AP3D1* or *B2M* as housekeeping genes for primers with annealing temperature at 60 °C (Hildyard and Wells 2014).

### Transcriptomic analysis

Total RNAs were extracted using NucleoSpin RNA Plus XS kit. RNA quality control was performed using Agilent 2100 Bioanalyser. Global gene expression was obtained using an Affymetrix GeneChip Human 8X60K chip. Data were normalized using LIMMA and controlled by principal component analysis before to compare the conditions with one-way ANOVA. Enrichment analysis was performed with DAVID gene ontology software. The transcriptome data are deposited at GEO as GSE229968.

### Generation of a library for ATAC-seq

The ATAC-seq libraries were generated using 50,000 cells from two technical replicates of 1 DMD sample. The samples were used as detailed in (Corces et al. 2017). In brief, nuclei were isolated and were incubated with a transposase solution (with a final concentration of transposase buffer 1X, 100 nM of transposase, 0.01% of digitonin, 0.1% of Tween-20 and PBS 0.33X) and incubated in a thermomixer with shaking at 1000 rpm for 30 min at 37°C. The mixture was cleaned using DNA Clean & Concentrator-TM-5 kit (Zymo #D4003). Next, transpose DNA was amplified by 12 cycles of PCR using SureSelectQXT Library Prep for WGS (Agilent #9684). The library quality and fragment size were quantified using an Agilent bioanalyzer 2100 before to be sequenced on Illumina HiSeq 4000 platform with paired-end sequencing. The ATACseq data are deposited at GEO as (undergoing).

### ATAC-seq data processing

The adapter sequences were trimmed using Cutadapt 2.6. Next, the reads were aligned to the reference hg38 genome using Bowtie 2 v2.3.4.1. The files were sorted and indexed with Samtools v1.10 and mitochondrial reads were removed. Peak calling was performed using MACS2 2.1.2 with the default q-value cut-off of 0.05 and keep-dup 1. Functional annotation of peaks was done with ChIPseeker 3.1.1.

### CUT&Tag

CUT&Tag was performed with mouse anti-HA antibodies (1:100, Genscript #A01244), rabbit anti-H3K4me3 antibodies (1:100, Sigma #04-745) or mouse anti-IgGs antibodies (1:100, Abcam #ab6708) using 7,500 cells as previously published (Li et al. 2021). In brief, after washes the cells were incubated with concanavalin A coated magnetic beads (Bangs Laboratories #BP531) for 15 min at RT. Bead-bound cells were incubated with the primary antibodies overnight at 4°C on a rotating platform. Primary antibodies were removed and mouse anti-IgGs antibodies (1:100, Abcam #ab6708) or rabbit anti-IgGs antibodies (1:100, Abcam #ab6708) were added for each condition and incubated for 1 h at RT. The cells were next incubated for 1 h at RT with pA-Tn5 adapter complex diluted at 1:250 before to proceed to the tagmentation for 1 h at 37 °C. The tagmentation was stopped by an overnight incubation at 37°C after addition of 18 µl of a solution composed of 0.5 M EDTA, 10% SDS and 10 mg/ml proteinase K. A phenol:chloroform:isoamyl DNA isolation was performed.

### Generation of the CUT&Tag library

To prepare libraries, 21 µl of DNA was mixed with 2 µl of universal i5 and i7 primers (Buenrostro et al. 2015) and 25 µl of NEBNext HiFi 2X PCR master mix (NEB #M0544). Samples were amplified in a thermocycler as follows: 72°C for 5 min, 98°C for 30 sec, 14 cycles of 98°C for 10 sec and 63°C for 30 sec and a final extension at 72°C for 5 min. Post PCR clean-up was performed with GeneJET PCR Purification kit (Thermo Scientific #K0701).

### Quantitative RT-PCR of CUT&Tag library

Each sample was tested in triplicate. Quantitative PCR was performed on CUT&RUN libraries using CFX96 Real-Time PCR Detection System (Bio Rad). The 10 µl final volume of reactive mixture contained 2 µl of diluted cDNA, 0.5 µl of CUT&TAG primer mixture (Table S5), 2.5 µl of water and 5 µl of LightCycler 480 SYBR Green I Master Kit (LifeSciences #04707518001). After initial denaturation of 2 min, the amplification was performed for 45 cycles of 95 °C for 10 sec, 60 °C for 5 sec and 72 °C for 10 sec (Hildyard and Wells 2014). The calculation of normalized relative quantity (NRQ) was performed by comparison of H2K9me3 and HA CUT&TAG samples to the IgG control samples.

### Statistics

All experiments were performed using at least 3 different donors (number of samples is given in the figure legends). Results are expressed using mean ± SEM. Statistics were performed using paired or unpaired t-tests or ANOVA and are given in the figure legends.

## Acknowledgments

This work was supported by FFCR (Fonds Franco-Canadien pour la Recherche), MITACS, Université Claude Bernard Lyon 1 to both BC and FJD labs, AFM-Telethon (Alliance MyoneurALP), Inserm, CNRS (to BC), and CIHR (FDN143330 to F.J.D.). We thanks the GENOMIC facility of Institut Cochin and the Cell Bank of Hopital Cochin.

## Author Contributions

Conceptualization: CG, ID, FJD, BC

Methodology, investigation: JM, MWG, HB, MM, AH, PN Analysis and Validation: JM, MWG, FJD, BC Resources: AH, PN, CG, ID

Writing and Visualization: JM, MWG, HB, MM, AH, PN, CG, ID, FJD, BC Supervision and funding acquisition: FJD, BC

